# Universal function of WRI1 involved in integrated regulation of multiple latex metabolism pathways in *Hevea brasiliensis* and *Taraxacum kok-saghyz*

**DOI:** 10.1101/2024.10.22.619728

**Authors:** Ruisheng Fan, Jiong wan, Wenfeng Yang, Fang Wei, Honghua Gao, Peng Qu, Huabo Du, Jian Qiu

**Affiliations:** Ministry of Agriculture and Rural Affairs Key Laboratory of Biology and Genetic Resources of Rubber Tree, Rubber Research Institute, Chinese Academy of Tropical Agricultural Sciences, Haikou, Hainan 571101, China; College of Tropical Crop Science, Yunnan Agricultural University, Puer, Yunnan 665000, China

## Abstract

*Hevea brasiliensis* and *Taraxacum kok-saghyz* are sources of nature rubber (NR) and recent research has focused on finding a method to increase NR quality and production. NR consists mainly of *cis*-1,4-polyisoprene, which is synthesized from sucrose by a series of reactions during carbon metabolism. WRINKLED1 (WRI1) is a transcription factor (TF) that coordinates many genes involved in carbon metabolism and lipid biosynthesis. Here, we isolated and characterized two orthologues of *WRI* in rubber-producing plants *H. brasiliensis*a and *T. kok-saghyz*, which are highly expressed in their latex. Subcellular localization and ectopic expression in *Arabidopsis thaliana* indicated that the TFs, *HbWRI1* and *TkWRI1*, are involved in lipid accumulation. Overexpression of *HbWRI1* and *TkWRI1* in *T. kok-saghyz* substantially enhanced NR quality and production, including dry rubber content, molecular weight, and the diameter of rubber particles in the latex. Conversely, they were decreased in *TkWRI1* repression. Furthermore, based on the activated expression in transgenic *T. kok-saghyz* latex and the existence of AW-box element in the promoter regions, 16 direct downstream genes of HbWRI1 and TkWRI1 were identified by a dual-luciferase reporter assay and yeast one-hybrid assay, and their products were responsible for both NR biosynthesis and lipid metabolism. These results reveal a regulatory module that HbWRI1 and TkWRI1 TFs can positively regulate NR quality and production in *T. kok-saghyz* latex by synergistically regulating the entire NR biosynthesis. This module is expected to be utilized to develop superior varieties to enhance NR quality and production.

## Introduction

In the present world, natural rubber (NR) is an indispensable industrial material because of its unique physical and chemical properties, including resilience, elasticity, abrasion resistance, and tensile strength. These properties make NR superior to synthetic rubber and NR has been widely used in the automotive, electronic, medical, and construction industries (Mantello et al., 2014; Rahman et al., 2013; Sakdapipanich, 2007). Unlike synthetic rubber, NR consists principally of polyisoprene, as well as 3–5% non-isoprene components, such as lipids, proteins, and metal ions. Polyisoprene is the basic carbon skeleton structure. Lipids and proteins act as branching points linking with polyisoprene to form a network and they confer unique crystallization behavior and tensile strength (Yamashita and Takahashi, 2020; Lehman et al., 2022). More than two thousand plant species produce NR and they are commonly called rubber-producing plants. *Hevea brasiliensis*, a tropical tree species, which native to the Amazon

Basin, and currently it is the only viable source of commercial NR (Chen et al., 2023; Chao et al., 2023). Recently, *H. brasiliensis* cultivation has been expanded to other tropical and subtropical regions and currently supplies approximately 14 million tons of NR annually. However, this is still insufficient for the sustained growth in NR demand.

*H. brasiliensis* cultivation is subject to a number of challenges including disease, such as South American leaf blight, high labor requirements, and a long production cycle, which makes supply unpredictable (Salehi et al., 2021). *Taraxacum kok-saghyz* grows in temperate regions and produces high-quality NR similar to that of *H. brasiliensis*, and it represents a promising alternative NR crop. However, poor productivity and low biomass limit its commercial application (Salehi et al., 2021; Yamashita and Takahashi, 2020). By genome and genetic transformation, *T. kok-saghyz* is used as a model plant species for studying rubber biosynthesis (Lin et al., 2022; Kuluev et al., 2023). To meet the ever-increasing demands for NR, an in-depth understanding of the biosynthesis and metabolic engineering of rubber-producing plants is necessary.

The latex from which NR is manufactured is produced in laticifers. The white or yellowish milky latex is secreted from the laticifers and harvested by tapping of the trunk bark periodical (Chao et al., 2015; Lestari et al., 2015; Tian et al., 2015). Polyisoprene, as the major component of latex, is derived from a continuous metabolic process, including isopentenyl pyrophosphate (IPP) biosynthesis and IPP polymerization. IPPs are produced primarily by the cytosolic mevalonate (MVA) and plastids methyl erythritol-4-phosphate (MEP) pathways using pyruvate and pyruvate-derived acetyl-CoA as a substrates (Yamashita and Takahashi, 2020). IPP polymerization is unique to latex metabolism and occurs in rubber particles (RPs), the specialized organelles in the latex, comprising a hydrophobic polyisoprene core surrounded by a half-unit phospholipid membrane and membrane proteins. Rubber transferase protein complex on the surface of RPs catalyze rubber chain extensions, and they including small rubber particle protein (SRPP), rubber elongation factor (REF), rubber transfer enzyme (HRT), HRT1-REF bridging protein, etc (Yamashita et al., 2016). Therefore, polyisoprene is a product of complex latex metabolism involved in pyruvate metabolism, isoprenoid biosynthesis, phospholipid synthesis, and RPs formation. Transcription factors (TFs) have been found to modulate multiple enzyme genes involved in polyisoprene biosynthesis in *H. brasiliensis*, such as *HbTGA1* (Guo et al., 2022), *HbWRKY27* (Qu et al., 2020), and *HbMYC2* (Deng et al., 2018). However, the coordinating regulation of multiple metabolic latex processes to produce polyisoprene still remains unclear.

Lipids, such as fatty acids (FAs), glycerides, and phospholipids are essential metabolites in plant cells, playing important roles in signal transduction, membrane biogenesis, energy storage, and stress responses (Xu et al., 2023; Yang and Benning, 2018). Previous studies in *H. brasiliensis* have shown that lipids are the most abundant non-rubber components in NR (over 3.0% of dry weight), and have been found to affect the NR properties (Liengprayoon et al., 2013). The lipid detailed compositions of latex and RPs have been described in both *H. brasiliensis* and *T. kok-saghyz*. Nine classes of phospholipids are abundant in the latex of both plant species, and most of them are produced from RPs (Bae et al., 2020). It has been speculated that phospholipids, which constitute the skeleton of RPs, play a key role in NR biosynthesis and RPs structure. However, the lipid biosynthesis in laticifers of both plants is little known. WRINKLED1 (WRI1) is the first TF found in *Arabidopsis thaliana* that is key to the regulation of lipid biosynthesis. The *AtWRI1* (At3g54320) mutant *wri1-1* seeds show an 80% reduction in triacylglycerol (TAG) accumulation and a five-fold increase in erucic acid, leading to delayed embryo elongation and wrinkled seeds (Baud et al., 2007; Cernac and Benning, 2004). The WRI1 encoding an APETALA2 ethylene responsive element-binding protein (AP2/EREBP) TF, involved in global regulation of lipid biosynthesis and carbon metabolism, was confined to high-expression in oil-enriched tissues of plants, such as seeds (Qiao et al., 2022).

In previous study, we found that two *WRI1* homologous gene (designated *HbWRI1* and *TkWRI1*) were abnormally highly-expressed in latex of *H. brasiliensis* and *T. kok-saghyz*. Laticifer is not a lipid rich tissue, however pyruvate is produced via glycolysis. It is possible that *WRI1* may turn on or boost the carbon flow in IPP biosynthesis and polymerization in laticifer tissues that provide a previously unknown regulatory mechanism of NR biosynthesis. In this study, we report that in the laticifers of *T. kok-saghyz*, HbWRI1 and TkWRI1 regulates production, RPs diameter, and molecular weight of NR by over-expression and RNAi-repression. We characterized a novel regulatory mechanism of WRI1 involved in polyisoprene biosynthesis integrating multiple latex metabolic processes, including carbon metabolism, lipid biosynthesis, and RP formation. Our research provides new insights into the regulation of latex metabolism, and could be significant in enhancing NR quality and production of rubber-producing crops.

## Results

### Enrichment of lipid and rubber biosynthesis in the latex transcriptome of *H.brasiliensis*

To gain insights into the molecular mechanisms underlying latex biosynthesis, transcriptomic profile of the latex and bark were investigated through RNA-Seq analysis from *H. brasiliensis*. According to cut-off criteria (|log_2_Fold Change| > 2 and adjust *P value* < 0.05), a total of 13,014 differentially expressed genes (DEGs) between latex and bark and 3,799 up-regulated DEGs were identified as tissue-specific expression genes in latex. Functional enrichment analyses of these genes by both GO and KEGG revealed that they were mainly enriched in the lipid biosynthetic process (GO:0008610), isoprenoid biosynthetic process (GO:0008299), terpenoid backbone biosynthesis (hbr00900), protein processing in endoplasmic reticulum (hbr04141), and glycerophospholipid metabolism (hbr00564). In addition, 9,215 DEGs were down-regulated in latex and were mainly enriched in the phenylpropanoid biosynthetic process (GO:0009699; hbr00940), plant-type primary cell wall biogenesis (GO:0009833), photosynthesis (hbr00195), and starch and sucrose metabolism (hbr00500). There have been reports that lipid metabolism is more abundant in the latex of other rubber yielding plants, including *Lactuca sativa* and *T. kok-saghyz*. These results suggest that lipid and isoprenoid biosynthesis could play a key role in the biosynthesis of latex.

As lipid biosynthesis is a dominant pathway in the latex, a *WRI1* homologous gene highly expressed in latex (log_2_Fold Change = 2.51) was observed from the gene set of the lipid biosynthetic process (GO:0008610) (Figure 1A). As anticipated for a *WRI1* homologous gene was also identified from the latex of *T. kok-saghyz*. The two *WRI1* homologous genes were designated as *HbWRI1* and *TkWRI1*, respectively. The qPCR results showed that *HbWRI1* and *TkWRI1* were detected in all the tested tissues of *H. brasiliensis* and *T. kok-saghyz*, with the highest abundance in latex (Figure 1, B and C). In addition to the latex, the expression level of *HbWRI1* in flowers and leaves was higher than in other organs (Figure 1B), and the expression level of *TkWRI1* in leaves was higher than that in roots and flowers (Figure 1C). The results indicate that *HbWRI1* and *TkWRI1* are associated with latex metabolism.

**Figure 1.**
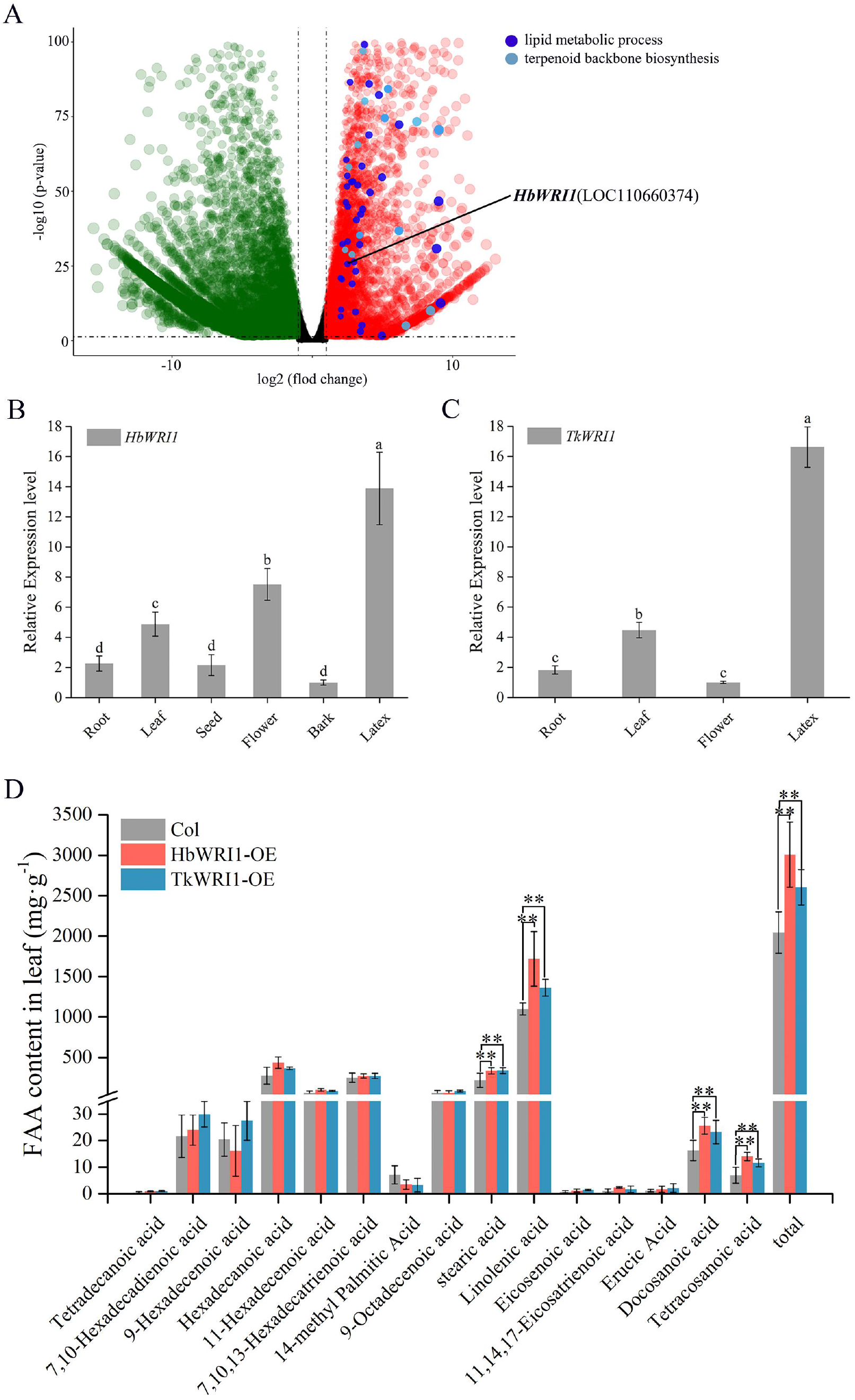
*HbWRI1* and *TkWRI1* regulate lipid biosynthesis. A, DEGs of latex and bark from *H. brasiliensis.* Red color circles represent latex up-regulated DEGs and green color circles represent bark up-regulated DEGs. B, *HbWRI1* organ-differential expression level in *H. brasiliensis.* C, *TkWRI1* organ-differential expression level in *T. kok-saghyz.* D, FA contents in HbWRI1 and TkWRI1 heterologous expression in leaves of *A. thaliana.* Date are presented as means ± SE. The Duncan test was performed to identify significant differences between groups. Asterisk indicates significant differences (*P* ≤ 0.05). Experiments were repeated three times.

### HbWRI1 and TkWRI1 are TFs that regulate lipid biosynthesis

The amino acid sequence relationships showed that two WRI1 homologous sequences from *H. brasiliensis* and *T. kok-saghyz* have a close relationship with WRI1 sequences from various plants (Figure S1). The subcellular localization in *Nicotiana benthamiana* leaf showed that HbWRI1-GFP (green fluorescent proteins) and TkWRI1-GFP fusion proteins were found in the nucleus (Figure S2), which suggests that HbWRI1 and TkWRI1 are imported into the nucleus to potentially function as TFs. To further investigate the functions of HbWRI1 and TkWRI1 in the regulation of lipid metabolism, *HbWRI1* and *TkWRI1* transgenic plants were obtained by heterologous expression in *A. thaliana*, and the FA composition and content in leaves were determined. Compared with the wild-type, the content of major FA, including stearic acid (C18:0, SA), linoleic acid (C18:2, LA), docosanoic acid (C22:0), and tetracosanoic acid (C24:0) in the leaves of transgenic HbWRI1 and TkWRI1 were significantly increased, resulting in a rise of total FA (Figure 1D). These results suggest that HbWRI1 and TkWRI1 possess TF characteristics and positively regulate the lipid biosynthesis.

### Latex metabolism and rubber production in transgenic *T.kok-saghyz*

To understand the potential function of HbWRI1 and TkWRI1 in laticifer cells, heterologous expression of *HbWRI1*, and over expression and RNAi of *TkWRI1*, in *T. kok-saghyz* were conducted driven by both CaMV35S promoter (abbrev 35) and laticifer-special REF promoter (abbrev Rp) (Epping et al., 2015). This generated three types of transgenic *T. kok-saghyz*: HbWRI1-h (35HbWRI1 -h and RpHbWRI1-h), TkWRI1-o (35TkWRI1-o and RpTkWRI1-o), and TkWRI1-i (35TkWRI1-i and RpTkWRI1-i). As confirmed by RT-qPCR (Table S1), *HbWRI1* was ectopically expressed in the latex of HbWRI1-h plants, and *TkWRI1* was over-expressed and repressed in TkWRI1-o and TkWRI1-i, respectively. There was no obvious difference in appearance in the transgenic and wild-type *T. kok-saghyz* (Figure 2A), but the expression level of objective genes differed substantially, along with various transgenic lines (Table S1). Five independent lines of each transgenic type were selected for comparison of relative latex content (RLC), dry rubber content (DRC), and FA content according to the expression level of corresponding genes (high in HbWRI1-h and TkWRI1-o, and low in TkWRI1-i).

**Figure 2.**
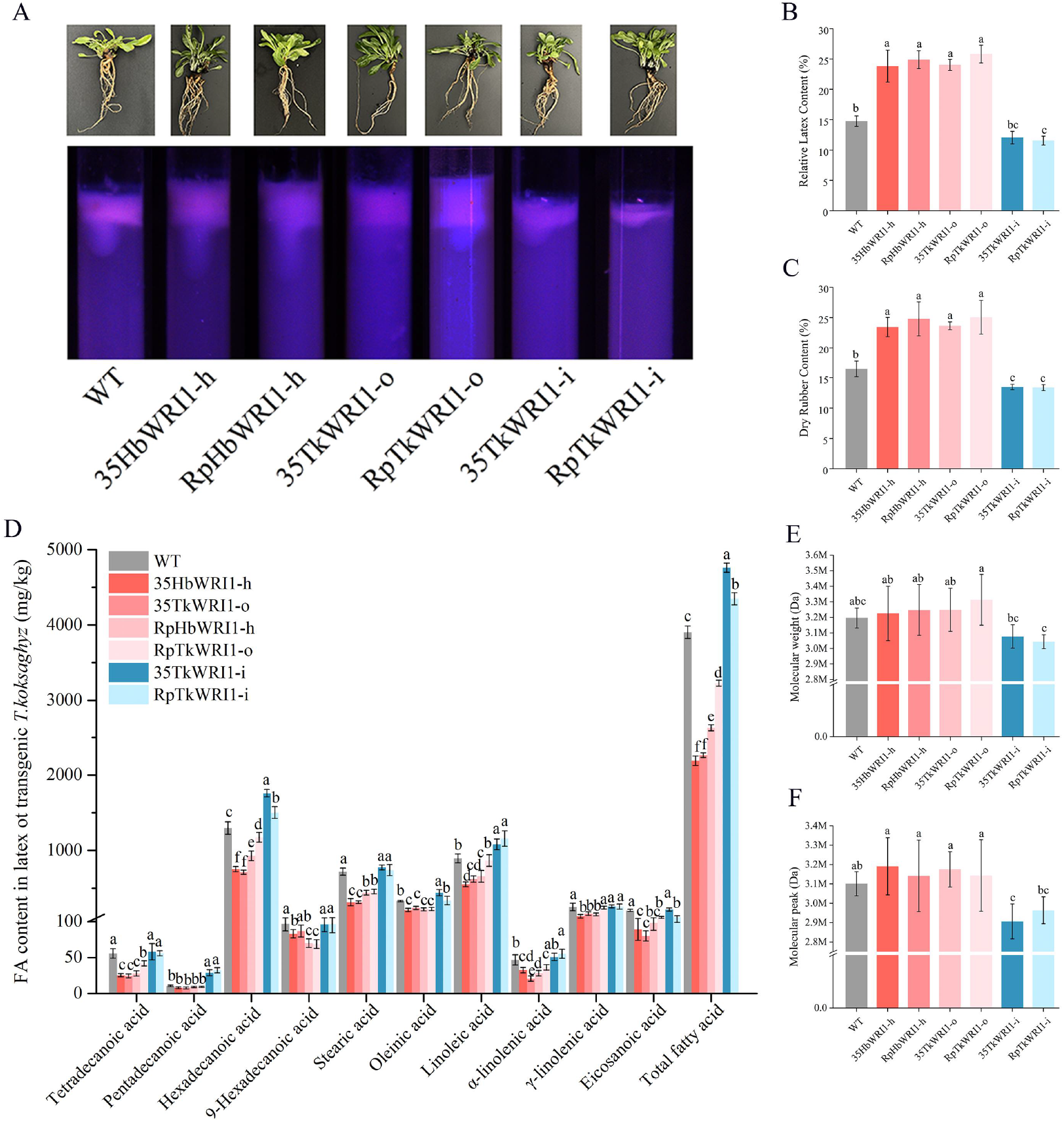
Characterization of latex in *WRI1* Transgenic *T. kok-saghyz*. A, Appearance of wild-type and transgenic *T. kok-saghyz*. B, The RLC of wild-type and transgenic *T. kok-saghyz*. C, The DRC of wild-type and transgenic *T. kok-saghyz*. D, FA content of wild-type and transgenic *T. kok-saghyz* latex. E, Mw of wild-type and transgenic *T. kok-saghyz*. F, Mp of wild-type and transgenic *T. kok-saghyz*. Each value and error bar in the graphs represent the mean of three biological replicates ± SD. The Duncan test was performed to identify significant differences between groups. Different letters indicated significant differences (*P* ≤ 0.05), Experiments were repeated three times.

The contents of RLC, DRC, and FA were altered in three types of transgenic plants compared to their wild-types. There was a significant increment of RLC (wild-type: 14.77%) in HbWRI1-h and TkWRI1-o plants (35HbWRI1-h: 23.82%; RpHbWRI1-h: 24.88%; TkWRI1-o: 24.03%; RpTkWRI1-o: 25.82%. *P*<0.05), and a small decrement in TkWRI1-i plants (35TkWRI1-i: 12.05%; RpTkWRI1-i: 11.56%. *P*<0.05) (Figure 2B), while the DRC (wild-type: 16.44%) was significantly enhanced in HbWRI1-h and TkWRI1-o plants (35HbWRI1-h: 23.44%; RpHbWRI1-h: 24.78%; TkWRI1-o: 23.64%; RpTkWRI1-o: 25.06%. *P*<0.05), and reduced in TkWRI1-i plants (35TkWRI1-i: 13.46%; RpTkWRI1 -i: 13.36%. *P*<0.05) (Figure 2C). The composition and total contents of FA were further determined in transgenic and wild-type *T. kok-saghyz*. Hexadecanoic acid (C16:0), stearic acid, oleic acid (C18:1), and linoleic acid were detected as the major FAs in latex. Both HbWRI1-h and TkWRI1-o plants accumulated significantly lower total FA, with reductions of most major FAs. By contrast, in TkWRI1 -i plants the total and major FA showed a strong increase compared to wild-type (Figure 2D). These results suggest that HbWRI1 and TkWRI1 are involved in the regulation of latex and lipid metabolism, and could be a positive regulation factor for rubber biosynthesis.

### Molecular weight and rubber particle size of NR in transgenic *T.kok-saghyz*

To understand the function of WRI1 in rubber biosynthesis, the average molecular weight (Mw) and peak average molecular weight (Mp) of latex were measured. A high molecular mass (more than 3×10^6^ Da) of rubber was detected in wild-type and transgenic plants by gel permeation chromatography (Figure S3). The Mw of latex was slightly increased in HbWRI1-h and TkWRI1-o, and decreased in TkWRI1-i plants compared with wild-type (Figure 2E). The variations of Mp in transgenic lines were similar to Mw; nevertheless, the decrease in TkWRI1-i plants was significant (*p*<0.05) (Figure 2F). The size of RPs and the molecular weights of latex were further examined. The average diameter of RPs from HbWRI1-h and TkWRI1-o plants ranged from 77.39 to 96.91 nm, which was significantly larger than that of the wild-type at 61.70 nm. While the average diameter of RPs from TkWRI1-i plants were 57.31 nm and 58.98 nm, both were smaller than wild-type (Figure 3A). Compared to wild-type, the number of large RPs in HbWRI1-h and TkWRI1-o plants and the accumulation of small RPs in TkWRI1-i plants were observed using transmission electron microscopy (TEM) (Figure 3, B-H). These results indicate that over-expression of *HbWRI1* and *TkWRI1* contribute to the accumulation of RPs and formation of high Mw rubber, but this was reversed for *TkWRI1* repression.

**Figure 3.**
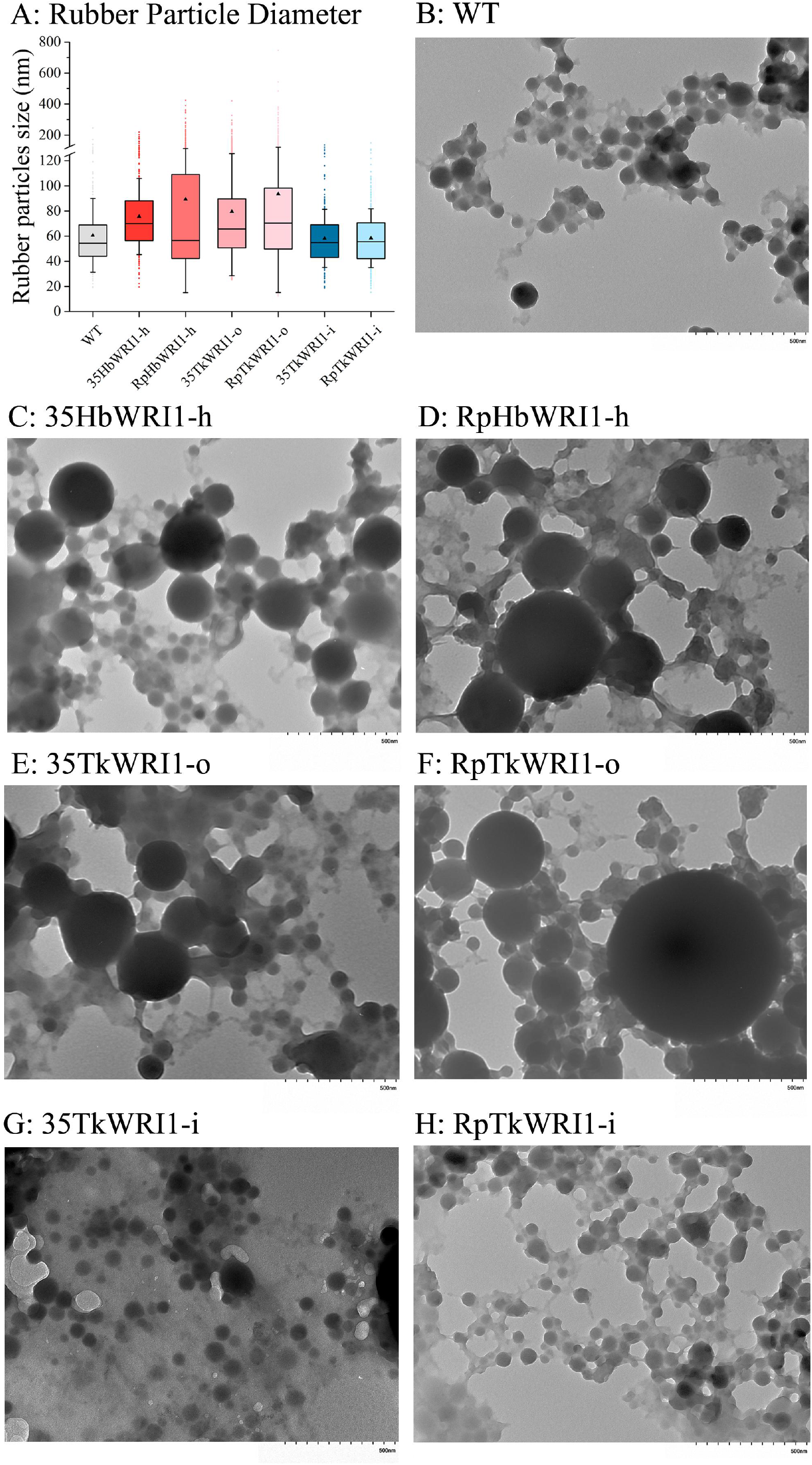
Characterization of RPs in *WRI1* Transgenic *T. kok-saghyz*. A, RPs diameter distribution of wild-type and transgenic *T. kok-saghyz.* B-H, TEM of latex from wild-type and transgenic *T. kok-saghyz.* Dark circle represents RPs. The horizontal line represents the median, the triangle represents the mean, the error line represents the ± SE, and the dot represents the outlier.

### Multiple pathways related to latex metabolism in transgenic *T. kok-saghyz*

A substantial effect on latex metabolism was observed in transgenic *T. kok-saghyz*. Transcript profiles of latex among wild-type and transgenic *T. kok-saghyz* were compared to identify downstream pathways related to WRI1 and elucidate further the regulation mechanism in latex metabolism. In *T. kok-saghyz*, the up-regulated DEGs in the TkWRI1-o and the down-regulated DEGs in the TkWRI1-i were identified as potential downstream genes of TkWRI1, while the up-regulated DEGs in HbWRI1-h were targets of *HbWRI1*. Compared with the wild-type, there were 1,252 DEGs up-regulated in TkWRI1-o and 366 DEGs down-regulated in TkWRI1-i, including *TkWRI1*. These genes were significantly enriched in 30 KEGG pathways and 16 KEGG pathways, respectively, including glycolysis/gluconeogenesis, carbon metabolism, pyruvate metabolism, and terpenoid backbone biosynthesis (Figure S4; Table S2). In HbWRI1-h, 875 up-regulated DEGs were identified, and most were significantly enriched in 29 KEGG pathways, including carbohydrate metabolism, glycerolipid metabolism, and sesquiterpenoid and triterpenoid biosynthesis (Figure S4; Table S2). A total of 312 DEGs known to be involved in carbohydrate metabolism, rubber biosynthesis, and fatty acid biosynthesis were identified as the potential downstream genes of TkWRI1 or HbWRI1 in latex. The main genes with significant differences are labeled on the corresponding metabolic pathways in Fig. 4 (Table S3 and S4). These results revealed that WRI1 acts as a positive regulator of multiple latex metabolism, playing crucial roles in controlling NR biosynthesis and latex production.

**Figure 4.**
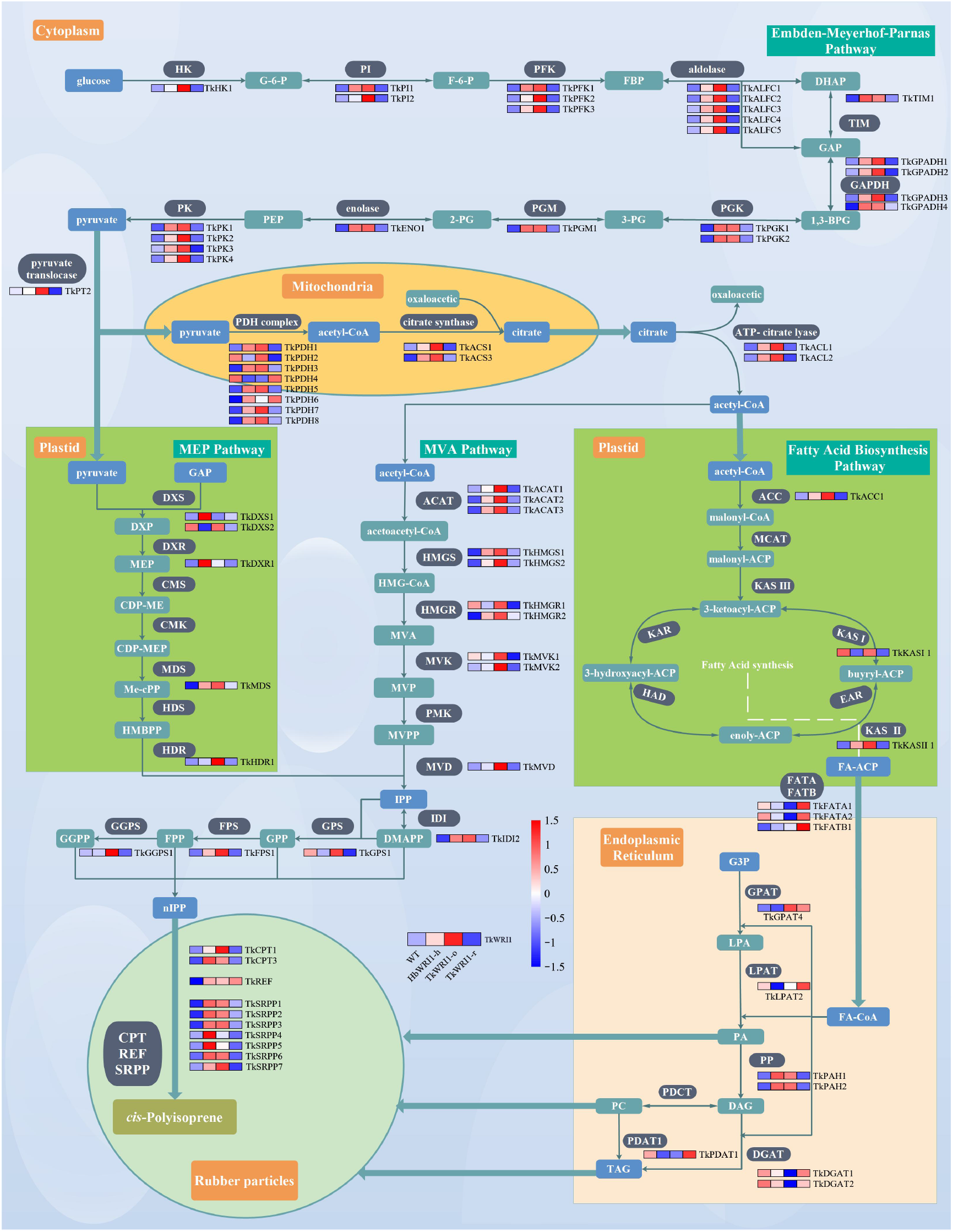
Biosynthesis of lipid and NR and the related isoprenoid pathway in transgenic *T. kok-saghyz.* Expression profiles of each enzyme gene in transgenic plants are represented by a heat map. All abbreviations are displayed in Table S4.

### WRI1 activates lipid metabolism and latex biosynthesis via binding AW-box of gene promoters

Plant WRI1 is known to activate genes via binding to the AW-box motif (CG[N]_7_CnAnG). Given the finding that WRI1 is involved in carbohydrate metabolism, FA, and rubber biosynthesis in latex, we wanted to verify that these conservative regulatory mechanisms are applicable to metabolic pathways in laticiferous cells, To do this, the potential targets of WRI1 were identified by the occurrence of AW-box. The interaction between two WRI1 and the promoters (Table S5) was further detected by dual-luciferase reporter and yeast one-hybrid (Y1H) assays. Almost a third of genes involved in the above metabolic pathways showed more than one AW box in their promoter, mainly enriched in sucrose metabolism, glycolysis, and FA biosynthesis, such as *PK*, *PDH*, *KAS*, and *KAR*. Notably, the AW-box motif is also present in the promoters of many genes related to rubber biosynthesis (Table S3), which could indicate that the AW-box is the binding site of WRI1 regulating multiple metabolic pathways. To test this hypothesis, the 1000-bp promoter of the selected genes containing AW-box were cloned from *H. brasiliensis* and *T. kok-saghyz* (Figure 5A). For *HbWRI1* as an effector, the firefly luciferase and renilla luciferase (LUC/REN) ratios significantly increased in the *HbPK2*, *HbPDH3*, *HbMVD1*, *HbGGPS1*, *HbKAR1*, *HbKASI 1*, and *HbLPAT1a* promoters compared to the negative control (empty reporter vector) (Figure 5B and S5). For *TkWRI1* as the effector, a significant increase in relative LUC/REN values was found in the *TkPHD1*, *TkPDH2*, *TkACAT1*, *TkHMGR1*, *TkDXS2*, *TkKASI 1*, *TkKASII 1*, *TkFATB1*, and *HbPDAT1* promoter compared to the negative control (Figure 5C and S5). Moreover, interaction among two WRI1 and promoters of 16 selected genes were further detected by Y1H assay using around 200 bp promoter containing AW-box. Yeast cell harboring WRI1 and the enzyme gene promoter survived on the selective medium. These results suggest that an interaction between WRI1 and AW-box of promoter did occur, and both HbWRI1 and TkWRI1 drove the enzyme gene transcription by binding to its AW-box in the promoter (Figure 5D and S6).

**Figure 5.**
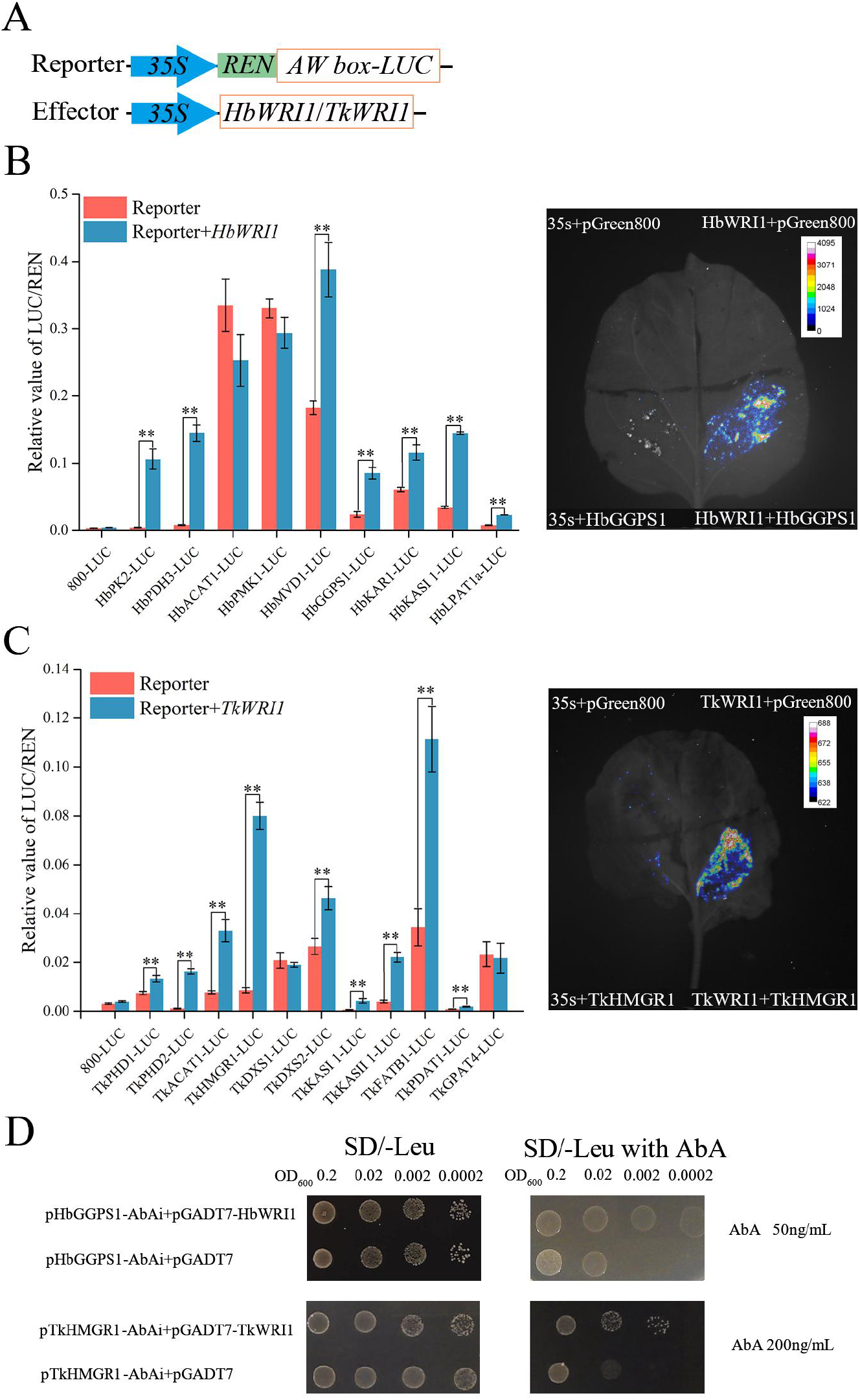
Validation of WRI1 transcription regulators in *vivo* and in *vitro*. A, Schematic representation of the effector and reporter plasmid. B, Transient transcriptional activity assays of HbWRI1 on enzyme gene promoters for dual-luciferase assays and *HbGGPS1 in vivo*. Other enzyme genes are displayed on Figure S5. C, Transient transcriptional activity assays of TkWRI1 on enzyme gene promoters for dual-luciferase assays and *TkHMGR1 in vivo*. Other enzyme genes are displayed on Figure S5. D, Transient transcriptional activity assays of HbWRI1 and TkWRI1 on AW-box of *HbGGPS1* and *TkHMGR1* promoters for Y1H assays. Other enzyme genes are displayed on Figure S6. Each value and error bar in the graphs represent the mean of three biological replicates ± SD. Asterisk indicates significant differences (*P* ≤ 0.05).

## Discussion

### The laticifer *WRI1* is functional in lipid biosynthesis

Previous studies showed that WRI1 has multiple functions in plants, including lipid biosynthesis, flowering, root development, seed development and stress responses (Baud et al., 2009; Cernac et al., 2006; Chen et al., 2018; Hao et al., 2017; Kong et al., 2017; Li et al., 2015; Marchive et al., 2014; Qu et al., 2012). These research data suggest that *WRI1* is involved in the global regulation of carbon metabolism. HbWRI1 and TkWRI1 proteins have two AP2/ERF functional domains, which specifically bind to the ethylene response element, GCC-BOX (Figure S2). They are linked together by 27 conserved amino acid sequences (Guo and Ecker, 2004; Sasak et al., 2007; Shinozaki and Yamaguchi-Shinozaki, 2000), which demonstrate the high conserved nature of *WRI1* across different species. Usually, highly conserved genes have similar functions, which suggests that the function of *HbWRI1* and *TkWRI1* may be similar to that in other plants. In the first AP2 domain, there is a VYL motif and 14-3-3 protein-binding motif. The VYL motif is a crucial component for WRI1 function. Point mutation of a single amino acid in VYL in *A. thaliana* can cause functional impairment of AtWRI1 protein (Ma et al., 2013). The 14-3-3 protein-binding motif contributes to AtWRI1 protein stability and transcriptional activity (Ma et al., 2016). The WRI1 contains three intrinsically disordered regions (IDR), which may be caused by differences between the N-terminal and C-terminal addition sequences of WRI1 during the evolution of different species (Tang et al., 2019). In addition, TkWRI1 also contains a PEST motif that can be phosphorylated in the IDR3 region, which contains a large number of phosphorylation sites; therefore the absence of this motif or mutation of the phosphorylation site may increase the stability of WRI1 protein (Ma et al., 2015; Yang et al., 2019). The PEST motif was not found in the IDR3 region of HbWRI1, indicating that HbWRI1 protein was more stable than TkWRI1.

Both *HbWRI1* and *TkWRI1* were both located in the nucleus (Figure 1C), indicating that *HbWRI1* and *TkWRI1* mainly play a regulatory role in the nucleus, which is consistent with the results of most plants *WRI1* as a TF, regulating glycolysis and lipid biosynthesis transcription (Baud et al., 2009). Heterologous expression of *HbWRI1* and *TkWRI1* in *A. thaliana* showed a rise of total FA content in leaves (Figure 1D), suggesting that they play roles in lipid biosynthesis. In general, *WRI1* should be expressed at high levels in plant organs with a higher lipid content, such as flowers, ovules and seeds. However, *HbWRI1* and *TkWRI1* expression levels were highest in latex (Figure 1, A and B), which has not been reported in previous studies. Considering that the lipid content in latex is approximately 3%, the ultra-high expression of *HbWRI1* and *TkWRI1* indicates that this TF may have other functions in latex. Given that the biosynthesis substrates of latex and lipid are the same, we reasonably speculate that *HbWRI1* and *TkWRI1* may play an important role in regulatory latex metabolism.

### *WRI1* is a positive regulator of NR biosynthesis by affecting latex and lipid metabolism

At present, no reports of the effect of WRI1 on latex metabolism were found, although the role of WRI1 on glycolysis and lipid metabolism has been extensively studied (Chen et al., 2020; Ruuska et al., 2002). *HbWRI1* and *TkWRI1* isolated from laticifer tissue showed a lipid metabolism function by heterologous expression in *A. thaliana* leaves (Figure 1D), but it remains unknown whether expression of WRI1s in different organs affect their functions. It would be more convincing to show the functions of HbWRI1 and TkWRI1 in the regulating latex metabolism if we could express or knock down their *WRI1* specifically in the laticifer of *H.brasiliensis* or *T. kok-saghyz* and find an increase or decrease of latex accumulation. Although an anti-oxidative stress superoxide dismutase gene was first inserted in transgenic *H.brasiliensis* in 2003 and several TFs have been transformed to *H.brasiliensis* successfully in recent years, the genetic transformation of *H.brasiliensis* is still a formidable challenge for gene function research (Jayashree et al., 2003; Lestari et al., 2018; Wang et al., 2023). Therefore, in this study *HbWRI1*, *TkWRI1*, and RNAi of *TkWRI1* were transferred to *T. kok-saghyz* (Knyazev et al., 2017; Qiu et al., 2014) to test their functions in laticifer. We observed that ectopic expressed *HbWRI1* and over expressed *TkWRI1* promoted rubber biosynthesis, which corresponded with the significant increase of RLC and DRC in *T. kok-saghyz* (Figure 2, B and C). Meanwhile, silencing *TkWRI1* repressed rubber biosynthesis.

Apart from the production of NR, transgenic *T. kok-saghyz* also has an impact on NR quality. Over expression *WRI1* in laticifer increased the molecular weight (Mw and Mp) of dry rubber (Figure 2, E and F) and particle diameter of RPs (Figure 3), but these were decreased in *TkWRI1* repression. In general, high molecular weight latex has better mechanical strength and elasticity because longer molecular chains are able to tangle and self-strengthen more effectively (Sakdapipanich et al., 2015). Molecular weight also affects the aging process of NR production and high molecular weight latex because of its more dense and uniform network structure, which is conducive to improving the final strength and durability of the product (Attanayake et al., 2018; Lei et al., 2018). For latex, large RPs have more non-rubber substances adsorbed on the surface, a thicker double electric layer and hydration film, a higher potential energy peak, and they are more stable; such a structure can provide better impact strength (Dai et al., 2013). Larger RPs can result in an increased latex viscosity and may be suitable for applications requiring higher filler content or specific flow characteristics (Sakdapipanich et al., 2015; Zhang et al., 2021). In summary, overexpression of *WRI1* in laticifer can improve the quality of NR in *T. kok-saghyz*.

In addition, we were surprised to find that over expression in *WRI1* significantly reduced the content of total and major FA in laticifer. By contrast, in *TkWRI1* repression the content of total and major FA were increased (Figure 2D). As a TF responsible for lipid biosynthesis, overexpression of *WRI1* in laticifer actually reduced the content of lipid in latex, which was contrary to previous conclusions observed in other plants (Baud et al., 2007; Huang et al., 2019; Liu et al., 2018; Vogel et al., 2019; Yang et al., 2019; Ye et al., 2018; Zhang et al., 2017). TThe decrease in lipid content of *WRI1* transgenic *T. kok-saghyz* lines may be due to several reasons. First, the laticifers were not the main site of lipid biosynthesis. Most of the enzyme genes involved in lipid biosynthesis were not highly expressed in latex. Second, acetyl-CoA is the common precursor of latex and lipid biosynthesis (Figure 4), aand rubber-producing plants adjust the amount of carbohydrate to meet metabolic demand for latex at the expense of lipid biosynthesis (Silpi et al., 2007).

### Regulation of *WRI1* TF at the molecular level

Based on current knowledge, several TFs have been identified that promote NR biosynthesis. HbTGA1 up-regulates the expression of multiple NR biosynthesis genes, such as *HbCPT8*, *HbHMGR2* and *HbSRPP2* (Guo et al., 2022). HbWRKY27, HblMYB19, and HblMYB19 positively regulate the expression of *HbFPS1* (Qu et al., 2020; Wang et al., 2017). HbMYC2 activates *HbFPS1* and *HbSRPP1* expression (Deng et al., 2018). Therefore, the identification of TFs involved in latex metabolism is necessary to understand the polyisoprene biosynthesis pathway in more detail, but also to apply them to improve NR both qualitatively and quantitatively. Based on the transcriptome of *WRI1* transgenic *T. kok-saghyz* latex, at least one gene of *TkHK*, *TkPI*, *TkPFK*, *TkALFC*, *TkGAPDH*, *TkPGM*, *TkENO*, *TkPK*, *TkPT*, *TkPDH*, *TkACS*, *TkACL*, *TkACAT*, *TkHMGR*, *TkMVK*, *TkMVD*, *TkDXR*, *TkCMK*, *TkMDS*, *TkHDR*, *TKGPS*, *TkFPS*, *TkGGPS*, *TkCPT*, and *TkSRPP* were up-regulated in HbWRI1-h and TkWRI1-o transgenic lines and down-regulated in TkWRI1-i transgenic lines (Table S3), respectively. These *WRI1*-regulated enzyme genes contained almost every key step of NR biosynthesis (Figure 4). These results indicate that *WRI1* activate enzyme genes of Embden-Meyerhof- Parnas (EMP), MVA, MEP, and NR polymerization pathways in *T. kok-saghyz* latex to synergistically regulate the entire NR biosynthesis. This echoes the new function of *WRI1* involved in NR biosynthesis which we previously proposed. In addition, we found that *TkWRI1* regulated some enzyme genes involved in early FA biosynthesis, such as *TkACC*, *TkMCAT*, *TkKAR*, *TkHAD*, *TkKASII*, and *TkKASIII* (Figure 5 and Table S3). This is the same as the previous studies conclusion (To et al., 2012; Yang et al., 2019).

Previous studies have found that AW-box is close to the transcription start site, which is easy to play its function and has a wide range of existence (Fukuda et al., 2013). The AtWRI1 activates genes involved in late EMP and early FA biosynthesis and binds to AW-box directly (Baud et al., 2009; Maeo et al., 2009; To et al., 2012). Similarly, the GmWRI1a can successfully bind to AW-box in soybean (Chen et al., 2018). According to transcriptome data, we selected nine gene promoters of *H. brasiliensis* and eleven gene promoters of *T. kok-saghyz* which have at least one AW-box in their promoters and contain each pathway involved in NR and FA biosynthesis (Table S5). The results of dual-luciferase reporter assay and Y1H indicated that HbWRI1 and TkWRI1 binds to the AW-box of seven and nine enzyme gene promoters (Figure S5 and S6), respectively. These results suggest that HbWRI1 and TkWRI1 activate genes involved in NR biosynthesis. These genes include *HbPK2*, *HbPDH3*, *HbMVD1*, *HbGGPS1*, *TkPHD1*, *TkPDH2*, *TkACAT1*, *TkHMGR1*, and *TkDXS2*, and they may help us to understand the regulatory mechanism of WRI1 involved in NR biosynthesis. In addition, not all *WRI1*-regulated enzyme gene promoters have an AW-box. We speculated that *WRI1* may indirectly regulate enzyme genes through other TFs or direct target genes, such as *HbMYC2-like1/2*, *MYB35-like*, and *GLB1* (Baud and Lepiniec, 2010; Deng et al., 2018; Wang et al., 2017).

## Conclusion

TFs that regulate NR biosynthesis improve NR quality in a relatively macroscopic way because TFs tend to regulate a range of gene expressions, which is closer to the collaborative process in the organism. This study is the first successful functional analysis of *WRI1* in transgenic *T. kok-saghyz* and it adds a new layer of regulation mechanistic detail to NR biosynthesis from latex. Overexpression of *WRI1* enhance production and quality of NR from latex. The evidence presented in our study shows that *WRI1* establish a key role for positive regulated NR biosynthesis by activating genes involved in pathways of EMP, MVA, and MEP, IPP polymerization, and early FA biosynthesis (Figure 6). Given that *WRI1* should have pleiotropic effects, further characterization will be needed to identify the target genes associated with NR biosynthesis. Knowledge of the details of this regulatory circuit can be used to design strategies to increase NR accumulation for biotechnological applications of *H. brasiliensis* in the future.

**Figure 6.**
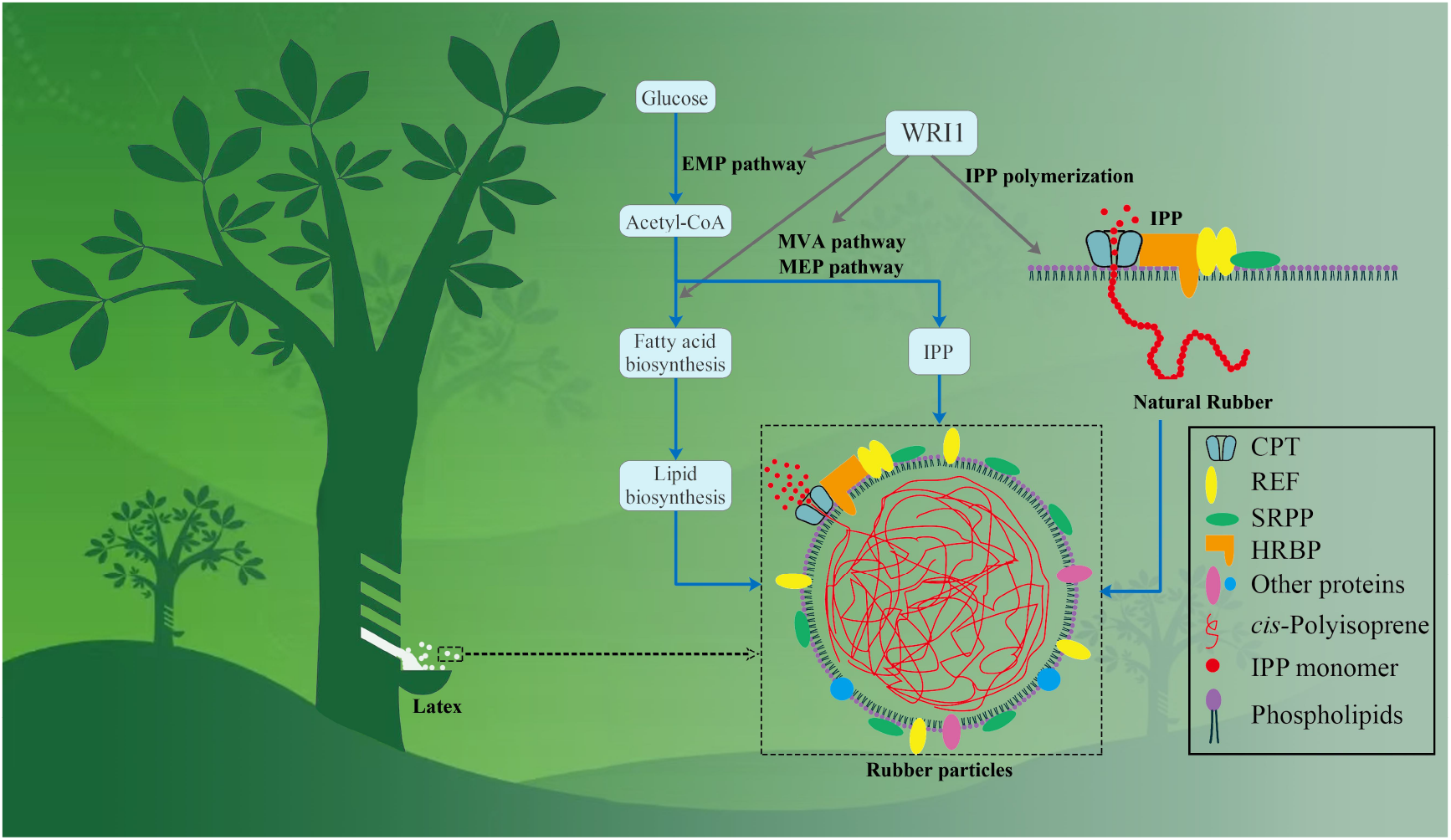
Regulatory network of WRI1s in NR biosynthesis and schematic diagram of RPs structure.

## Materials and methods

### Plant material and cultivation

The latex was tapped from 10-year old *H. brasiliensis* CATAS 73397, which was grown at experimental plantation of the Rubber Research Institute (Danzhou, Hainan, China). The latex harvesting system was performed by half spiral tapping (S/2) once every four days (d4), without ethephon application. According to the standard harvesting system, latex and barks were collected in liquid nitrogen and stored at −81℃ for later use. Latex was discarded in the last 30 s after tapping. Leaves, flowers, seeds, and roots were collected in liquid nitrogen and stored at −81℃ extraction extraction of total RNA.

T. *kok-saghyz* wild-type was provided by Yang et al. (2021) and planted in the germplasm nursery of the Rubber Research Institute (Haikou, Hainan, China), and transgenic lines were transformed and regenerated in July 2021. In April 2022, the tissue culture plantlets were transferred from culture flasks into the soil in pots (12 cm × 12 cm × 15.5 cm) and cultivated at 24℃ with a 16 h photoperiod. After 5 months, the latex from each transgenic plant was harvested by picking the roots with a needle. The latex of *T. kok-saghyz* was used in the experiment immediately after needling roots. Leaves, flowers, and roots were collected and stored at −81℃ until extraction of total RNA.

### Total RNA isolation and quantitative real-time PCR (qRT-PCR)

Total RNA was isolated from the tissue of *H. brasiliensis* and *T. kok-saghyz* using the EasyPure Plant RNA Kit (TranGen, Beijing, China) according to the manufacture instructions. Total RNA from the latex of *T. kok-saghyz* was isolated using the E.Z.N.A^®^ MicroElute Total RNA Kit (OMEGA, Norcross, USA) according to the manufacture instructions. The purity and concentration of RNA was checked using NanoDrop 2000 (Thermo Scientific, Waltham, USA). The cDNA was synthesized using the PrimeScript^TM^ Fast RT reagent Kit with gDNA Eraser (TaKaRa, Dalian, China). The qRT-PCR reactions were conducted in a final volume of 20 μL, using the TB GREEN^®^ Premix Ex Taq^TM^ II Fast qPCR (TaKaRa, Dalian, China) according to the user manual, and performed by qRT-PCR (CFX96, BIO-RAD, USA). Relative expression levels were normalized by the expression level of the internal control gene *HbYLS8* and *TkGAPDH*, calculated using the 2^−ΔΔCt^ method. The expression of each gene was performed in triplicate. All primers in this assay were listed in Table S6.

### Transcriptome analysis of *H. brasiliensis* and WRI1 transgenic *T. kok-saghyz*

The cDNA library construction and *de novo* transcriptome sequencing were performed by Novogene Co. Ltd (Tianjin, China). After generating the library, an Agilent 2100 Bioanalyzer was used to detect the inserted fragment scope of the library, and the ABI StepOnePlus real-time PCR System was used to quantify the library concentration. After quality inspection, the constructed cDNA libraries were quantified on the Illumina NovaSeq 6000 HiSeq platform. Raw data were first filtered by an internal software to remove the reads containing adapters. Reads containing more than 5% ambiguous poly-N and low-quality reads (reads with Phred score less than 10 accounted for more than 20% of the total base as low-quality reads) to generate clean reads. Data volume and quality value of clean reads were calculated, and all clean reads were *de novo* assembled by Trinity v. 2.0.6 software. The assembled transcript was clustered, and redundancies were removed by Tgicl and assessed with BUSCO.

Assembled unigenes were annotated using Pfam databases. The Gene Ontology (GO) annotation was performed using Blast2GO and SwissProt annotation results. Clean reads were aligned to the assembled reference sequence by Bowtie 2. The expression levels of unigenes and transcripts were calculated by RSEM and measured by FPKM. DESeq2 was used for DEGs analysis by the negative binomial distribution principle, according to Anders et al. (2010). Unigenes with |fold change| ≥ 2.00 and *p*-values ≤ 0.05 were defined as DEGs. According to the results of DEGs testing, DEGs were clustered by the pheatMap function in R software for hierarchical clustering analysis. According to the Kyoto Encyclopedia of Genes (KEGG) annotation results and the official classification, we classified the differentially expressed genes in KEGG pathways and used the phyper function in the R software to conduct enrichment analysis.

### Amplification HbWRI1 and TKWRI1 cDNA full-length sequences and bioinformatics analysis

The full-length cDNA of latex *HbWRI1* and *TkWRI1* were amplified using the degenerate primers *HbWRI1-F*, *HbWRI1-R*, *TkWRI1-F*, and *TkWRI1-R* and inserted into the clone vector T-vector pMD19 using T4 DNA Ligase (TaKaRa, Dalian, China). The final sequence construction was validated by Tsingke Biotech sequencing (Guangzhou, China). The open reading frame (ORF) of *HbWRI1* and *TkWRI1* sequence were discovered by the ORFfinder on NCBI and the different *WRI1* orthologs from other species were carried out via BLASTp on NCBI. The amino acid sequences alignment of *HbWRI1*, *TkWRI1* and *WRI1* orthologs from other species were analyzed by DNAMAN v6 software.

### Construction of *WRI1* recombinant plasmid

Subcellular localization recombinant plasmids were construction by CloneExpress Ultra One Step Cloning Kit V2 (Vazyme, Nanjing, China), the ORFs of *HbWRI1* and *TkWRI1* without stop codons were cloned into the N-terminal of the plasmid pCAMBIA1300-EGFP at the *Bam*H I site to construct *HbWRI1-EGFP* and *TkWRI1-EGFP*, respectively. Expression recombinant plasmids, transgenic for were construction by Gateway clone method. First, the ORFs of *HbWRI1* and *TkWRI1* were cloned into pGWC at the *Ahd* I site to construct *HbWRI1-pGWC* and *TkWRI1-pGWC*, respectively, using homology recombination protocols as the intermediate vector. Next, the expression vector pMDC32 (Curtis and Grossniklaus, 2003) was modified by replacing the Ref promoter (Epping et al., 2015), inserting a GUS tag, and constructing a hairpin structure. Then, the intermediate vector was attached to the expression vector using LR recombination protocols. Finally, the OE recombinant plasmids 35-*HbWRI1-*h, 35-*TkWRI1-*o, Rp-*HbWRI1-*h, and Rp-*TkWRI1-*o and RE (shRNA) recombinant plasmids 35-*TkWRI1-*r and Rp-*TkWRI1-*r were constructed using a GateV^TM^ LR Cloning Kit (Clone Smarter, USA).

### Subcellular localization of *HbWRI1* and *TKWRI1* in *N. benthamiana*

The subcellular localization recombinant plasmids were transferred into *Agrobacterium tumefaciens* GV3101 (pSoup-p19) (AngYu, Shanghai, China) (Nie et al., 2022). The localization vector MIEL1- mCherry was co-transformed with each combination as a nuclear marker (Miaoling, Wuhan, China). Transient transformation in *N. benthamiana* leaves epidermal cells as previously described (Liang et al., 2023) and modified to harbor the empty pCAMBIA1300-EGFP vector as a negative control. The fluorescence was observed under a confocal laser scanning microscope (Zeiss LMS800, German) with a 40× objective. The GFP excitation was performed using a 488 nm solid state laser and detected at 500–540 nm. The mCherry excitation was performed using a 552 nm solid state laser and detected at 560–620 nm. The images were post-processed using the Zeiss Zen-Blue-Lite software (Version 3.7).

### Heterologous expression of *HbWRI1* and *TKWRI1* in *A. thaliana*

The over expression recombinant plasmids were transferred into *A. tumefaciens* GV3101 (AngYu, Shanghai, China) harboring the empty pCAMBIA1303 vector as a negative control. *A. thaliana* was transfected by the floral dip method as previously described (Clough and Bent, 1998). Selection and detection of transgenic *A. thaliana* as previously described (Fan et al., 2019).

### Over expression and repress expression of *HbWRI1* and *TKWRI1* in *T. kok-saghyz*

Transformation in tissue of *T. kok-saghyz* was performed as previously described (Laffaldano et al., 2016), with slight modifications. In brief, roots were cut 1 cm from the wild-type *T. kok-saghyz* of 12 to 16-week old plants and immersed in a suspension of *A. tumefaciens* AGL1 (AngYu, Shanghai, China) carrying over expression and repress expression recombinant plasmids for 8 minutes under vacuum conditions. After 2–4 days, they were co-cultured in the dark, and root fragments were then transferred to a primary culture medium: solid Murashige & Skoog (MS) medium with 0.2 mg L^-1^ 3-indoleacetic acid (IAA), 0.5 mg L^-1^ kinetin (KT), 6 mg L^-1^ hygromycin (HYG) and 300 mg L^-1^ timentin (TIM). After 4 weeks of culture, regenerated callus was transferred to a subculture medium: solid MS with 0.1 mg L^-1^ IAA, 1.0 mg L^-1^ KT, 3 mg L^-1^ HYG and 150 mg L^-1^ TIM. They were cultured for a further 6–8 weeks and verified by GUS. Transgenic adventitious buds were selected and transferred to rooting medium: solid half-strength MS with 0.05 mg L^-1^ IAA, 10 mg L^-1^ HYG and 300 mg L^-1^ Tim. After further cultured on rooting medium for 8–12 weeks and verified by Genomic DNA PCR the transgene events were validated in selected tissue culture plantlets. Transgenic plantlets with thicker taproots were selected and they were then transferred to soil.

### FA analysis of *A. thaliana* leaves and *T. kok-saghyz* latex by GC-MS

FA analysis of *A. thaliana* leaves were performed as described previously (Fan et al., 2021). In brief, lyophilized leaves (20–30 mg) were transesterified directly in a glass tube by addition of 1mL n-hexane and 0.5mL of freshly prepared 5%(v/v) KOH-methanol. After reaction at 95°C for 2 h extracted the FA methyl ester (FAME) and analyzed FAME extracts by GC-MS (GC: Shimadzu UPLC LC-30A; MS: AB Sciex TripleTOF^®^ 6600). The FA contents were calculated by drawing the standard curve of FA external standard (37 kinds of non-esterified FA). The data were corrected for the fraction of the sample analyzed and normalized to the sample “dry weights” to produce data in the units mg/g.

For *T. kok-saghyz*, 20 μL latex and 10 μL whole FA internal standard were placed in a glass tube, 4 mL dichloromethane:methanol:water (2:1:0.8) solution was added and shaken. After centrifugation the supernatant was removed, and 2 mL dichloromethane was added to extract twice. Subsequently, the supernatant was combined and blow dried with nitrogen. The mixture was dissolved with 1 mL isopropanol and 0.5mL KOH-methanol. After methyl ester reaction, the FAME were extracted and analyzed by GC-MS. The analyzed method as before described with the units mg/kg.

### Analysis of RLC, DRC, and Mw in *T. kok-saghyz* latex

For RLC, 2 μL latex was collected by needling *T. kok-saghyz* roots. The latex was then transferred to 30 μL solution buffer (350 mM sorbitol, 100 mM Tris-HCl, pH<7.8). The latex-solution buffer-mixture was then transferred to a capillary tube and centrifuged for 5 min at 6000 g. The rubber phase was fractionated by centrifugation and observed under a fluorescent stereo microscope (LeiCa, M250FA, German). Latex volume was calculated by measuring the length of the rubber phase by image J software using the line measurement feature.

For DRC, 10 μL latex of *T. kok-saghyz* and 40 μL solution buffer were placed in a PCR tube and centrifuged at 18,000 g, 4°C for 20 min to fractionate the rubber phase. The rubber phase was transferred to a new centrifugal tube after which 10 μL acetic acid was added to solidify the rubber phase and air dried overnight. The DRC was obtained by the weighing method. The dry rubber was dissolved in tetrahydrofuran and the Mw distribution of rubber was analyzed by a gel permeation chromatography system (Waters, 1515 GPC, USA).

### TEM imaging and measurement of RPs diameter

TEM imaging of RPs was performed as previously described (Singh et al., 2003) and modified. In brief, 5 μL latex of *T. kok-saghyz* and 200 μL solution buffer were placed in a PCR tube and centrifuged at 18,000 g, 4°C for 20 min to fractionate the rubber phase. The rubber phase and part of the C-serum phase were transferred to a new centrifuge tube. Isotonic solution (250 mM sucrose, 100 mM Tris-HCl, pH=7.8) was added to the rinsed rubber and fixed to 200 μL. Six percent 200 μL glutaraldehyde solution (50mM sodium dimethylarsenate, 1% tannic acid) was added to fixed RPs for 2 h, and centrifuged to removed the supernatant. RPs were rinsed in isotonic solution three times, and 1% osmic acid was then added to the RPs at room temperature for 1 h. Isotonic solution was used to rinse RPs three times, after which they were centrifuged and collected. The RPs were fully suspended in 50 μL double distilled water. One drop of the suspended mixture on the copper net was volatilized and dried overnight. The coated sample was transferred to the TEM system (Hitachi, HT-7700, JAP). The sample was viewed at 80 kV, 8.0 μA with magnification at 20.0 k. RPs were visually identified and individual diameters of particles were measured in TEM images by Image J software using the line measurement feature.

### Downstream transcriptional regulatory genes of *WRI1* verified by dual-luciferase

According to the results of transgenic *T. kok-saghyz* transcriptome sequencing, *WRI1* coexpressed genes were screened. Protein and promoter sequences were extracted from the *H.brasiliensis* (Liu et al., 2020) and *T. kok-saghyz* genomes (Lin et al., 2018). Promoter sequences containing were used for MEME-ChIP to predict the binding elements of *WR1*. The target gene promoters were cloned into pGreenII800-LUC vector at the *Hind* III site to construct the report vector. The containing WRI1 protein overexpression recombinant plasmid and reporter vector (2:1) were co-transformed into *N. benthamiana* leaves by the *Agrobacterium*-mediated method, harboring the empty overexpression, and pGreenII800-LUC vector was used as a negative control. The fluorescence values of LUC and REN were detected by a dual-luciferase reporter gene assay kit (Zeye, Shanghai, China), and the LUC/REN ratio in each sample was calculated. The visual fluorescence of *N. benthamiana* leaves were visually identified using an Optical *in vivo* Imagine System (Photometrics, Lumazone pylon2048B, USA) and images were manipulated by Image J software.

### Transcriptional activity analysis of *WRI1* in Y1H system

Y1H system was performed as Y1HGold-pAbAi Yeast One-Hybrid interaction proving kit (Coolaber, Beijing, China). In brief, gene promoter elements were cloned into pAbAi vector at the *Hin*d III and *Xho* I sites to pBait-AbAi as bait vector. *HbWRI1* and *TkWRI1* were cloned into pGADT7 vector at the *Bam*H I and *Xho* I sites to AD-HbWRI1 and AD-TkWRI1 as prey vector. First, the self-activation and AbA resistance were tested using the bait vectors and the empty pGADT7 vector transformed into Y1H Gold yeast, and the AbA screening concentration was determined. Then, co-transforming bait and prey vectors validated *WRI1* interactions with promoter elements.

## Statistical analysis

Data were subjected to One-way ANOVA using SPSS and the Duncan test were performed to identify groups. Differences between treatments were considered to be significant at *P*<0.05. Each experiments were performed in triplicate.

## Accession numbers

The *T. kok-saghyz* transcriptome raw sequence data reported in this paper have been deposited in the Genome Sequence Archive (GSA) in National Genomics Data Center, database under CRA017701 that are publicly accessible at https://ngdc.cncb.ac.cn/gsa. The *H.brasiliensis* transcriptome raw data in FASTQ format from the present study can be accessed in the NCBI Sequence Read Archive (SRA) database under PRJNA1105020.

## Acknowledgments

We thank International Science Editing (http://www.internationalscienceediting.com) for editing this manuscript.

## Data availability

The data underlying this article are available in the article and in its online supplementary material.

## Author contributions

JQ designed the research, review & editing; RF and JW performed the experiments and analysis; RF analyzed data and wrote the manuscript; WY, HG, FW and PQ conducted data validation, and HD helped improve the manuscript.

## Supplementary data

The following materials are available in the online version of this article.

**Supplementary Figure S1.** DEGs of latex and bark from *H. brasiliensis.* Red color circles represent latex up-regulated DEGs and green color circles represent bark up-regulated DEGs.

**Supplementary Figure S2.** The Subcellular localization of *WRI1* in *N. benthamiana* leaf epidermal cells.

**Supplementary Figure S3.** The molecular weight with nonlinear curve fit.

**Supplementary Figure S4.** The significant GO pathway of transgenic *T. kok-saghyz* transcriptome.

**Supplementary Figure S5.** Transient transcriptional activity assays of HbWRI1 and TkWRI1 on all enzyme gene promoters for dual-luciferase assays and *in vivo*.

**Supplementary Figure S6.** The results of Y1H assays.

**Supplementary Table S1.** Information of transgenic *T. kok-saghyz* all compare.

**Supplementary Table S2.** Transcriptome data of transgenic *T. kok-saghyz* in KEGG pathway.

**Supplementary Table S3.** Selected downstream genes of WRI1 in biosynthesis lipid, NR and the related isoprenoid pathway in transgenic *T. kok-saghyz* and *H. brasiliensis*.

**Supplementary Table S4.** All abbreviations in Figure 4.

**Supplementary Table S5.** Selected enzyme genes to validated WRI transcriptional activity.

**Supplementary Table S6.** All primer in this assay.

**Supplementary Data Set 1.** The data of selected enzyme gene promoter sequences.

